# Augmenting Recombinant Antibody Production in HEK293E Cells: Optimising Transfection and Culture Parameters

**DOI:** 10.1101/2021.11.04.467283

**Authors:** Zealyn Shi-Lin Heng, Joshua Yi Yeo, Darius Wen-Shuo Koh, Samuel Ken-En Gan, Wei-Li Ling

## Abstract

**Background:** Optimising recombinant antibody production is important for cost-effective therapeutics and diagnostics. With impact on commercialisation, higher productivity beyond laboratory scales is highly sought, where efficient production can also accelerate antibody characterisations and investigations.

**Methods:** Investigating HEK293E cells for mammalian antibody production, various transfection and culture parameters were systematically analysed for antibody light chain production before evaluating them for whole antibody production. Transfection parameters investigated include seeding cell density, the concentration of the transfection reagent and DNA, complexation time, temperature, and volume, as well as culture parameters such as medium replacement, serum deprivation, use of cell maintenance antibiotic, incubation temperature, medium volume, post-transfection harvest day and common nutrient supplements.

**Results:** Using 2 mL adherent HEK293E cell culture transfections with 25 kDa linear Polyethylenimine in the most optimised parameters, we demonstrated a ~2-fold production increase for light chain alone and for whole antibody production reaching 536 and 49 μg respectively in a cost-effective manner. With the addition of peptone, κ light chain increased by ~4-fold to 1032 μg while whole antibody increased to a lesser extent by ~2.5-fold to 51 μg, with benefits potentially for antibodies limited by their light chains in production.

**Conclusions:** Our optimised findings show promise for a more efficient and convenient antibody production method through transfection and culture optimisations that can be incorporated to scale up processes and with potential transferability to other mammalian-based recombinant protein production using HEK293E cells.

**Statement of Significance:** Recombinant antibody production is crucial for antibody research and development. Systematically investigating transfection and culture parameters such as PEI/DNA concentrations, complexation time, volume, and temperature, supplements, etc., we demonstrated a ~4-fold light chain alone production increase to 1032 μg and a 2.5-fold whole antibody production increase to 51 μg from 2 mL transfections.

## INTRODUCTION

The efficient production of recombinant monoclonal antibodies underpins their functional and structural investigations (1) such as toxicology and drug formulation studies (2) and thereby, success of both therapeutic and diagnostic antibodies development. With a direct impact on production costs during development, optimisations of the already expensive mammalian production of recombinant antibodies are highly desired.

Currently, whole recombinant antibodies are predominantly produced by mammalian cells, given the need for complex folding, assembly and post-translational modifications to generate functionally active proteins (3, 4). Typically, the protein genes of interest are first transfected into mammalian cells, followed by the selection and generation of stable cell lines expressing the gene to produce the protein of interest (2). As this process is often tedious and laborious, the transient gene expression method on easily transfected cell lines is typically used for research scale investigations. One of the most widely used is the Human Embryonic Kidney 293 (HEK293) family of cell lines with several genetic variants developed to enhance protein expressions such as the HEK293E expressing the Epstein-Barr Virus Nuclear Antigen-1 (5), utilised in this study, and HEK293FT and HEK293T expressing the SV40 T-antigen (6). Given the scalability of HEK293 (3, 7, 8), they are suitable for producing a wide range of recombinant proteins including whole viral particles (9).

Given that transient recombinant protein production depends on a myriad of transfection and culture parameters, optimising these parameters directly influences the scalability and efficiency of recombinant protein production. Transfection parameters include seeding cell density (10), transfection reagent concentration (11), DNA concentration (12), complexation time, temperature, and volume (12), together with culture parameters such as medium replacements (7, 13), serum deprivation (14), use of cell line maintenance antibiotic (15), incubation temperature (16, 17), culture volume, harvesting time (18–20), and nutrient supplementation (21–23). Three main strategies are commonly used to optimise these parameters: the reductionist “one factor at a time” approach (24), stepwise optimisation from one condition to another (25), and the design of experiment approach (10), of which we took the single isolated factor approach.

In this study, the key drivers were determined by the “one factor at a time” approach and subsequently combined to maximise antibody production (24). We optimised individual parameters using Pertuzumab (Perjeta®) antibody light chains (Variable κ-1 or Vκ1), given that light chains are pre-requisites for whole antibody production (26, 27) and to mitigate the lopsided effect of dual plasmid transfections. This was followed by validating the combined parameters using recombinant whole Pertuzumab antibodies produced using two separate variable kappa (Vκ)-1 and variable heavy (VH)-3 plasmids. Based on our previous study where we generated Pertuzumab variants of VH family 1 to 7 and Vκ1 to 6 frameworks via complementarity determining regions (CDRs) grafting (28, 29), we analysed light chains known to limit antibody production i.e. the Vκ5 grafted Pertuzumab light chain as a pairing partner.

The final combination of optimised parameters for whole antibody production would complement our previous findings for production that discusses the role of heavy chain constant (30), Vκ-VH pairings (29), amino acid usage with leader selection (28) in a holistic manner (1, 31) to significantly boost antibody production for antibody-based therapeutics and diagnostics.

## MATERIALS & METHODS

### Plasmid DNA

Pertuzumab light chain of Vκ1 region and heavy chain IgG1 of the VH3 region genes, including their CDR-grafted V-region variants, designated Vκ2-6 and VH1, 2, 4-7 were ligated into pTT5 plasmid (YouBio, Cat no. VT2202) and prepared as previously described (28–30, 32, 33). Trastuzumab plasmid variants in Supplementary Figure 1B were prepared similarly. Optimisation of light chain production was performed by single plasmid transfection of Pertuzumab light chain Vκ1 prior to optimising whole Vκ1|VH3 antibody production by dual plasmid transfection with its corresponding heavy chain VH3. To investigate if the optimised parameters can boost production of antibodies limited by light chains, Pertuzumab Vκ5 was paired with VH3 for whole Vκ5|VH3 antibody production. Comparisons of culture incubation temperatures performed using other antibody variants in large flasks are shown in Supplementary Figure 1.

### Cell culture conditions

HEK293E adherent cells were routinely cultured in DMEM medium (Gibco, Cat no. 11965-084), supplemented with 10 % FBS (Gibco, Cat no. 10270-106) or 10 % Ultra Low IgG FBS (Gibco, Cat no. 16250078), 1 mM Sodium Pyruvate (Gibco, Cat no. 11360-070), 100 units/mL penicillin, 100 μg/mL Streptomycin (Nacalai tesque, Cat no. 09367-34), G418 (Roche, Cat no. 04727894001) and incubated at 37 °C in 5 % CO_2_.

### Transfection Parameters

Linear 25 kDa Polyethylenimine (PEI) (Polysciences, Cat no. 23966-1) at pH 7.0 was used in the transfections. 2 mL and 40 mL scale transfections of HEK293E cells were performed in 6-well plates (Costar, Cat no. 3516) and T175 flasks (Greiner Bio-one, Cat no. 660175) respectively, with at least three independent replicates. The conditions tested for the transient gene expression in 6-well plate format included: Cell seeding densities from 1 - 10 × 10^5^ cells/mL; PEI concentrations from 1 - 10 μg/mL; DNA concentrations from 0.25 - 2 μg/mL; PEI-DNA complexation time from 5 - 120 minutes; PEI-DNA complexation temperatures of 18 - 45 °C; PEI-DNA complexation in 1 - 25 % serum free medium (SFM) of total 2 mL volume; PEI-DNA complexation in light or dark environments; culture medium replacements; serum deprivation before transfection; culture with or without G418 antibiotic; culture incubation temperature at 32 or 37 °C; culture volume from 2 - 5 mL; post-transfection harvest of 3 - 21 days. Experimental conditions were compared against control parameters of: Seeding density at 2 × 10^5^ cells/mL; DNA concentration at 1 μg/mL; PEI concentration at 2 μg/mL; PEI-DNA complexation time at 30 minutes in 18 °C; with 10 % SFM of total 2 mL volume, performed in the dark and added into cultures without G418; with the cultures grown at 37 °C and harvesting made on day 14 post-transfection, as were performed routinely in previous work (28–30, 34).

For cell culture conditions, 10 % FBS DMEM was used for light chain production, while 10 % Ultra Low IgG FBS DMEM was used for whole antibody IgG production. The culture conditions used for Supplementary Figure 1 were scaled up proportionately from the 6-well cultures for use at T175 flask scale transfections.

### Supplement Preparation and Usage

The nutrient supplements tested were: OmniPur® Casamino acids (CA) (EMD Chemicals, Cat no. 65072-00-6), Casein from Bovine Milk (CS) (Sigma-Aldrich, Cat no. C3400), Bacto™ Peptone (PT) (Becton, Dickinson and Company, Cat no. 21167), Skim Milk (SM) (Millipore, Cat no. 70166), Tryptone (TT) (Bio basic, Cat no. TG217 (G211)), Bacto™ Yeast Extract (YE) (Becton, Dickinson and Company, Cat no. 212750) and Impact Essential Amino Acids (EA) (MYPROTEΓN™)(28). Supplements were dissolved in water or Phosphate Buffer Saline (PBS), filtered using 0.22 μm filters and stored at 4 °C before use.

The supplements were added post-transfection of Pertuzumab light chain cultures on day 7 at 1 mg/mL final concentration. Further optimisations with peptone were performed at 1, 2, 4 and 6 mg/mL, added on days 1 – 10. Subsequently, the combined optimised parameters of 1) without nutrient supplementation; 2) peptone at 1 mg/mL; 3) essential amino acids at 3.5 mg/mL (28) or 4) both were tested for Pertuzumab whole antibody production (Vκ1|VH3) and light chain limiting variant (Vκ5|VH3) with the supplementations (conditions 2, 3, and 4) added on day 7 post-transfection.

### Recombinant Antibody Harvest and Titre Quantification

Cell culture supernatants were harvested on various days stipulated by the experimental conditions and at the end of 14 days, and stored at −20 °C before further testing. In the small-scale 2 mL experiment set-ups where evaporation could be a potential confounding factor, total protein production was calculated from the concentration and remaining volumes measured by a micropipette at the point of harvest. The light chain and whole antibody quantification of the cell culture supernatants were performed using Octet Red 96 instrument as previously described (28–30), with Protein L (PpL) and G (SpG) biosensors (ForteBio, Cat no. 18-5085 and 18-5082) respectively.

### Data Analysis

The data were analysed and plotted using Microsoft Excel 365 and GraphPad Prism Version 9.0.0. Protein titre or total protein production data were normalised against the control and expressed as a percentage with the control set at 100 %. One-way ANOVA with Dunnett’s comparison test against the control was used when examining three or more different parameters, while a two-tailed unpaired parametric t-test was performed when comparing two parameters.

## RESULTS

### Optimising Transfection Parameters

#### Cell seeding densities

Cell seeding densities ranging from 1 - 10 × 10^5^ cells/mL in a 6-well plate were evaluated with respect to their impact on Pertuzumab light chain production and normalised against that of the control condition of 2 × 10^5^ cells/mL set at 100 % in **Figure 1A**. The one-way ANOVA showed no significant differences (*p* > 0.05) in Pertuzumab light chain production across the seeding densities tested despite a decreasing trend above 6 × 10^5^ cells/mL, where production at 10 × 10^5^ cells/mL was only 41 % (70 μg/mL) that of the control (177 μg/mL). The average titre data with accompanying standard error are shown in Supplementary Table 1.

**Figure 1:**
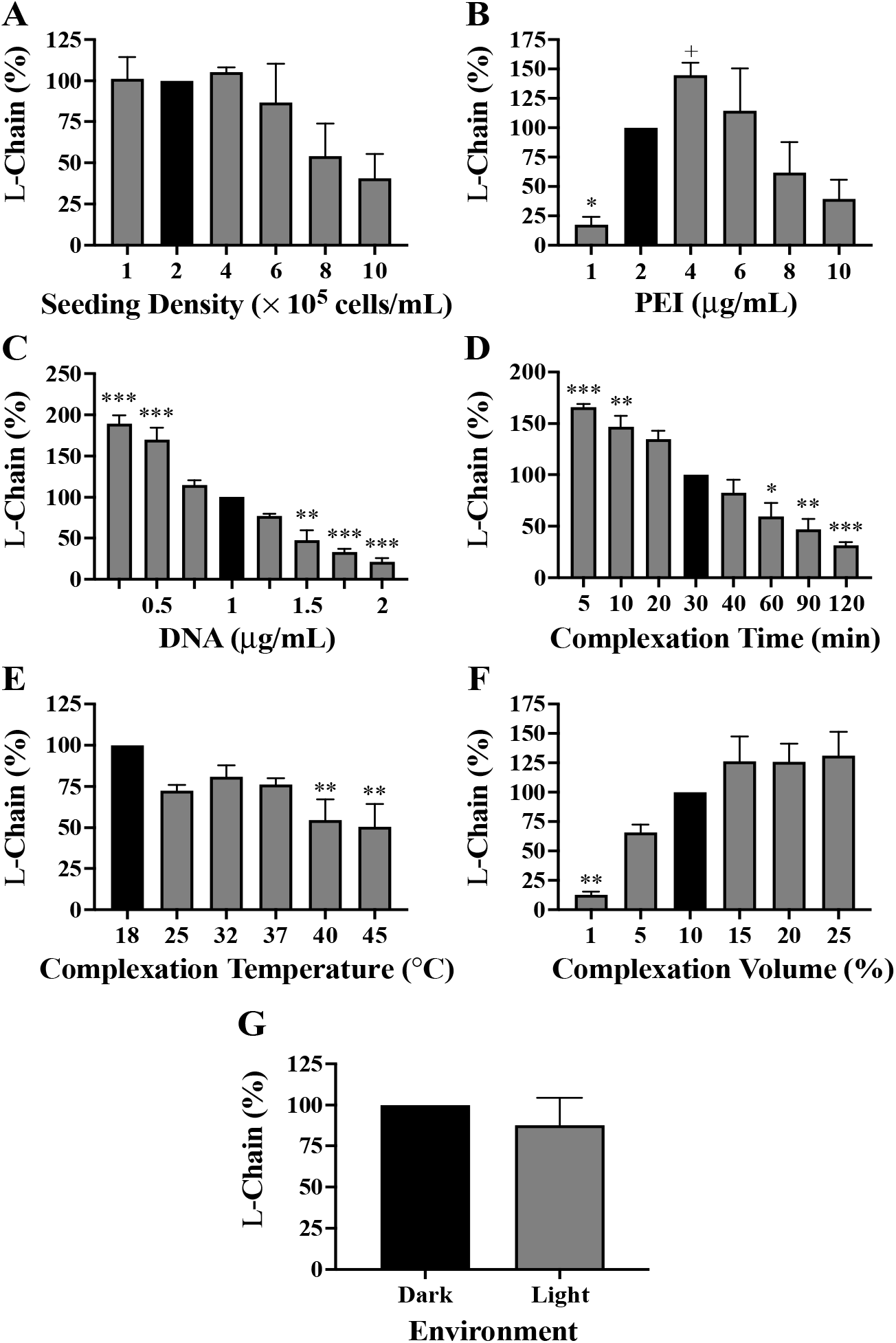
Pertuzumab light chain productions in 2 mL transfections from optimising the transfection parameters of (A) Seeding density; (B) PEI concentration; (C) DNA concentration; (D) PEI-DNA complexation time (E); PEI-DNA complexation temperature; (F) PEI-DNA complexation volume; (G) PEI-DNA complexation in light and dark environments. Black coloured bars represent the controls to which the experimental conditions were compared for statistical analysis. *, **, *** denote significance at *p* < 0.05, *p* < 0.01 and *p* < 0.001, respectively from one-way ANOVA Dunnett’s post-hoc test, while ^+^ denotes significance at *p* < 0.05 using t-test analysis. (The experiments were carried out in at least three independent replicates with titres and standard errors provided in Supplementary Table 1.)

#### PEI and DNA concentrations

PEI concentrations ranging from 1 - 10 μg/mL in 6-well plate transfections were tested for Pertuzumab light chain production and normalised against that of the control condition of 2 μg/mL set at 100 % in **Figure 1B**. The results showed a curvilinear relationship between PEI concentration and light chain production peaking at 145 % (206 μg/mL) for 4 μg/mL of PEI that was significant (*p* < 0.05) with t-test but not the Dunnett’s test against the control (140 μg/mL). PEI concentrations lower or higher than 4 μg/mL negatively impacted light chain production with 1 μg/mL having the lowest production of 17 % (22 μg/mL) significant at *p* < 0.05 when analysed with Dunnett’s test. Concentrations lower than 4 μg/mL also had lower production rates despite not being significant by Dunnett’s test.

DNA concentrations ranging from 0.25 - 2 μg/mL, at 0.25 μg/mL intervals in 6-well plate transfections, were evaluated for Pertuzumab light chain production and normalised against that of the control condition of 1 μg/mL set at 100 % in **Figure 1C**. An inverse relationship between DNA concentration and light chain production was observed with the lowest and highest DNA concentration (0.25 and 2 μg/mL) corresponding to the best and poorest light chain production at 190 % (399 μg/mL) and 21 % (64 μg/mL) respectively that were significant at *p* < 0.05 when analysed with Dunnett’s test against the control (208 μg/mL).

#### PEI and DNA complexation

PEI-DNA complexation time from 5 - 120 minutes in 6-well plate transfections was evaluated with respect to Pertuzumab light chain production and normalised against that of the control condition of 30 minutes, set at 100 % in **Figure 1D**. An inverse relationship between complexation time and light chain production showed the shortest and longest incubation time (5 and 120 minutes) corresponding to the highest and lowest production (166 % and 32 %, equivalent to a titre of 271 and 54 μg/mL respectively), which were both significant at *p* < 0.05 when analysed with Dunnett’s test against the control (165 μg/mL) production.

PEI-DNA complexation temperature from 18 to 45 °C in 6-well plate transfections was evaluated with respect to Pertuzumab light chain production and normalised against that of the control condition of 18 °C, set at 100 % in **Figure 1E**. An inverse relationship where the decline in light chain production was found with increasing temperatures. Dunnett’s test against the control for 25, 32, and 37 °C showed no significant decrease (*p* > 0.05) despite slight reductions. However, significant decreases (*p* < 0.05) were observed for 40 and 45 °C, at 54 % (105 μg/mL) and 50 % (98 μg/mL) of the control (164 μg/mL) production respectively.

PEI-DNA complexation volume using SFM from 1 to 25 % of the final total culture medium in 6-well plate transfections was evaluated with respect to Pertuzumab light chain production and normalised against that of the control condition of 10 % final cell culture volume (20) set at 100 % in **Figure 1F**. PEI-DNA complexation medium volume was found to increase light chain production with the lowest 1 % SFM yielding the lowest production of 12.7 % (30 μg/mL, *p* < 0.05) increasing to the highest production of 131 % (273 μg/mL) at 25 % SFM (*p* > 0.05). Nonetheless, production was observed to peak at 15 % transfection volume plateauing at higher complexation volumes.

To study photodegradation assumptions of transfection complexes (35), we performed PEI-DNA complexation incubation for light chain using the 6-well plate format under the biosafety cabinet (BSC) light and without (in very dim external room light). There was no significant difference (*p* > 0.05) in antibody light chain production between the two conditions (**Figure 1G**) by t-test.

### Optimising Culture Conditions

#### Culture medium replacements

Fresh medium replacement before or after transfection, as well as both before and after, were tested for Pertuzumab light chain using the 6-well plates and normalised against that of the control condition of no medium replacement set at 100 % in **Figure 2A**. Pertuzumab light chain production was found to increase by 36 % and 9 % (reaching 320 and 250 μg/mL) for before, and both before and after transfection, respectively. A slight reduction of 7 % to 235 μg/mL was found for a medium replacement after transfection when compared to the control (226 μg/mL), but the one-way ANOVA showed that these are not significant (*p* > 0.05). The average titre data with accompanying standard error are shown in Supplementary Table 1.

**Figure 2:**
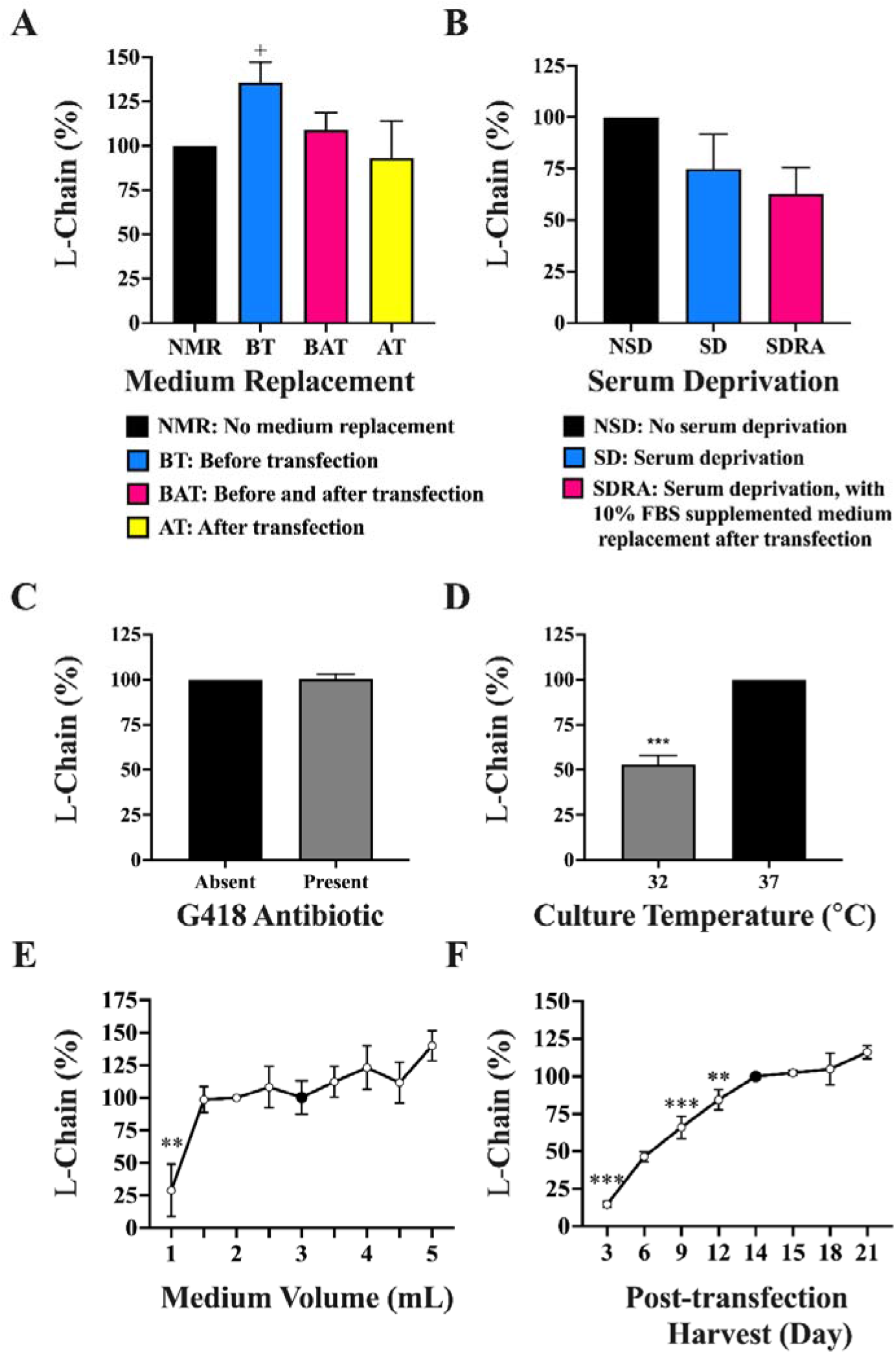
Pertuzumab light chain production in 2 mL transfections from optimising the culture parameters with respect to A) Medium replacement timing; (B) Serum deprivation; (C) Culturing with G418 antibiotic; (D) Culture temperature; (E) Culture medium volume (F) Post-transfection harvest day. Black coloured bars in A-D and black coloured dots in E and F represent the controls to which the respective experimental conditions were compared against for statistical analysis. *, **, *** denote significance at *p* < 0.05, *p* < 0.01 and *p* < 0.001, respectively from one-way ANOVA Dunnett’s post-hoc test. ^+^ denotes significance at *p* < 0.05 using t-test analysis. (The experiments were carried out in at least three independent replicates with light chain titres for A-D found in Supplementary Table 1 while the total protein for E and F can be found in Supplementary Table 2.)

We next studied serum deprivation by seeding cells in 10% FBS supplemented or serum free medium a day before transfection in 6-well plate format to test for Pertuzumab light chain production. An additional condition of seeding cells in serum free medium a day before transfection followed by replacement with 10% FBS supplemented medium a day after transfection was included. The serum deprivation condition data was normalised against that of no serum deprivation set at 100 % in **Figure 2B**. We found the prolonged and short-term serum deprivation to result in a non-significant (*p* > 0.05) 25 % and 37 % decrease in light chain production using Dunnett’s test against the control.

#### Culture with G418 antibiotics

Studying the effect of cell culture antibiotics on Pertuzumab light chain production in 6-well plates, we did not find G418 to have significant effects (*p* > 0.05) on the light chain production by t-test (**Figure 2C**).

#### Culture temperature

Culture incubation temperatures at 32 and 37 °C for Pertuzumab light chain 6-well, transfections were shown with t-tests to reveal a significant production decrease at 32 °C (*p* < 0.05). 2 mL cultures grown in 32 °C resulted in only 53 % (120 μg) of the light chain produced at 37 °C, set at 100 % (231 μg) (**Figure 2D**). The average total light chain data with standard error are shown in Supplementary Table 2. Further testing with a full panel of whole recombinant Pertuzumab and Trastuzumab variants (VH1-7 and Vκ1-6 combinations) transfected in T175 flasks did not show a significant difference (*p* > 0.05) between the two temperatures (Supplementary Figure 1).

#### Culture medium volumes

Medium volumes varying from 2 - 5 mL in 6-well plate transfections were evaluated for Pertuzumab light chain production and normalised against that of the control condition of 2 mL set at 100 % in **Figure 2E**. The Dunnett’s test showed a significantly lower difference (*p* < 0.05) when using a culture volume of 1 mL, producing 148 μg of total light chain compared to the control of 300 μg from a 2 mL culture. While there seems to be a gradual increase in light chain production for higher volumes used, with the highest medium volume of 5 mL showing a rise of 40 % (reaching 413 μg), it was not significantly different (*p* > 0.05) from the control.

#### Culture supernatant harvest day

Post-transfection harvest of culture supernatants across 3 – 21 days from Pertuzumab light chain 6-well plate transfections were normalised against that of the control condition of 14 days set at 100 % in **Figure 2F**. We found a direct relationship between the day of supernatant harvesting and light chain production where total light chain from 2 mL cultures increased from 81 μg on day 3 to plateau at 522 μg from day 14 (Supplementary Table 2). With analysis by Dunnett’s test against harvest Day 14, the control, there was an expected significantly lower (*p* < 0.05) light chain production on harvest days 3, 6, and 9. No significant increase (*p* > 0.05) was detected after day 14.

#### Nutrient Supplements

A variety of culture supplements commonly used to improve recombinant protein production were evaluated for Pertuzumab light chains and normalised with respect to the control condition of no supplement (NS) set at 100 % in **Figure 3A**. Adding the supplements to post-transfected cultures on day 7 at a final concentration of 1 mg/mL, the one-way ANOVA showed no significant difference (*p* < 0.05) across the supplements tested. Given that peptone is the most common supplement of mammalian recombinant protein production (25, 36–38), further investigations were performed using a two-tailed unpaired t-test against the non-supplemented control, revealing that it significantly boosted (*p* < 0.05) light chain production by 25 % (**Figure 3A**) to 351 μg in an initial 2 mL culture (Supplementary Table 2). A similar t-test was performed for yeast extract showing no significant (*p* > 0.05) increases. Further optimisations of concentration and timing of peptone addition showed that 1 mg/mL, added on post-transfection days 5, 6, or 7, gave the highest boost to light chain production (**Figure 3B**). Single supplementation of peptone at 1 mg/mL alone on day 1 or day 7 was found sufficient to enhance light chain production (**data not shown**), showing that repeated addition of low concentrations of peptone was not necessary when a mid-point culture addition on day 7 was sufficient.

**Figure 3:**
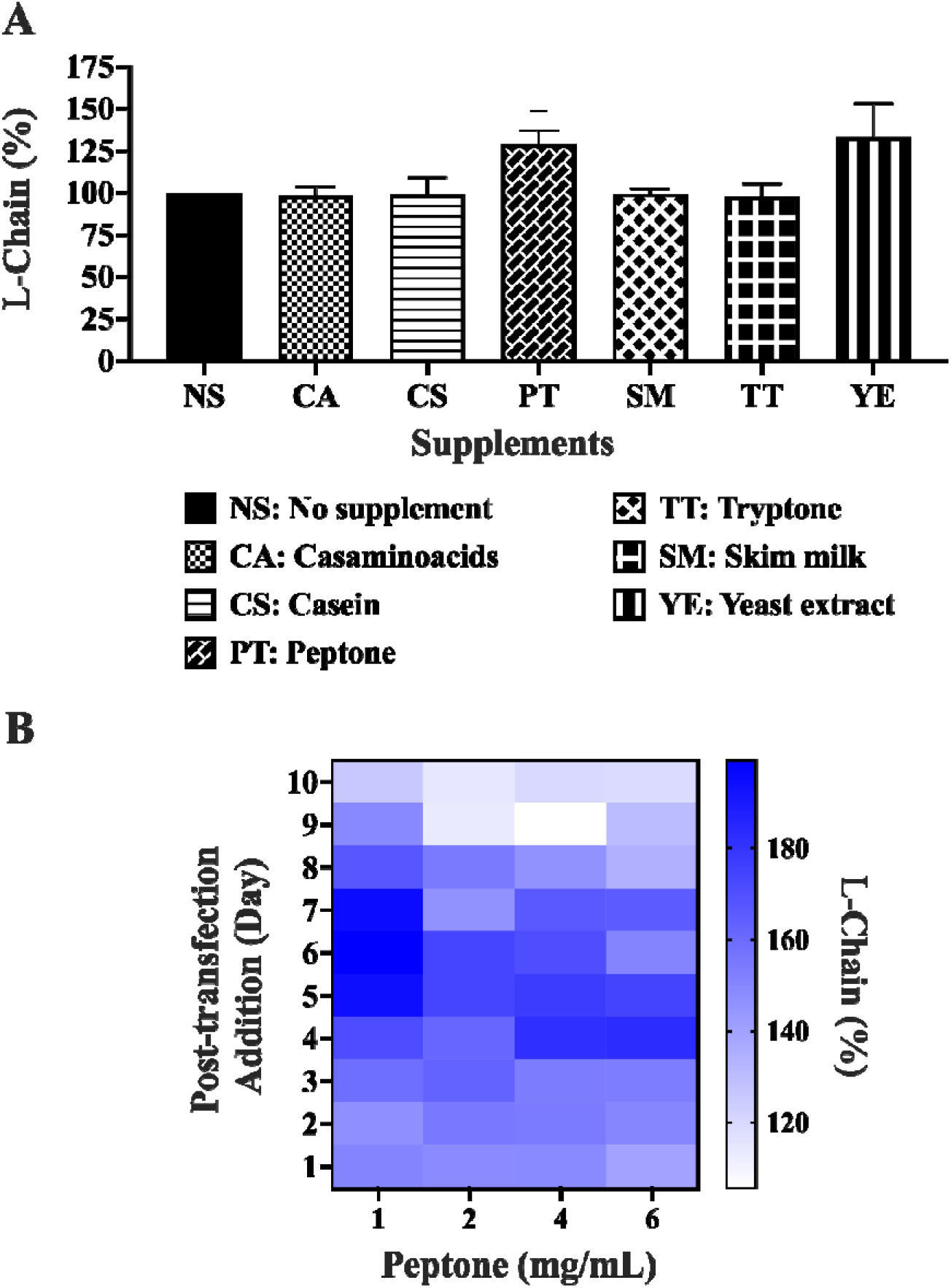
Antibody light chain production in 2 mL transfections with (A) nutrient supplements and (B) Heat map of one-time additions of different peptone concentrations. Black coloured bars in A represent the control to which the respective supplements were compared for statistical analysis. ^+^, denotes significance at *p* < 0.05 using t-test analysis. (The experiments were carried out in at least three independent replicates where the light chain total protein values are provided in Supplementary Table 2).

### Combination of Transfection and Culture Parameters

From 2 mL culture optimisations of the individual transfection and culture parameters in **Table 1**, the combined parameters for Pertuzumab light chain production led to a 217 % significant increase (*p* < 0.05) to 536 μg. This increase was further enhanced ~2-fold to 452 % reaching 1032 μg after peptone supplementation (**Figure 4A** and Supplementary Table 3). Validating this on whole Pertuzumab antibody production in 2 mL cultures, a significant increase (*p* < 0.05) of 236 % to 49 μg was elicited from the control of 22 μg. With peptone supplementation, a subtle increase of 252 % was observed, reaching 51 μg (Supplementary Table 3).

**Table 1:** Control and optimised parameters for transient recombinant production in 2 mL, 6-well plate cultures.

| Parameters | Control | Optimal |
| --- | --- | --- |
| Cell Density (cells/mL) | $2 \times 10^5$ | $2 \times 10^5$ |
| PEI Concentration (µg/mL) | 2 | 4 |
| Plasmid DNA Concentration (µg/mL) | 1 | 0.25 |
| Complexation Time (min) | 30 | 5 |
| Complexation Temperature (°C) | 18 | 18 |
| Complexation in SFM<br>(% of transfection volume) | 10 | 10 |
| Culture Medium Volume (mL) | 2 | 2 |
| Post-transfection Harvest (day) | 14 | 14 |
| Culture Incubation Temperature (°C) | 37 | 37 |
| Supplements (Post-transfection) | - | 1 mg/mL Peptone on Day 7 |

**Figure 4:**
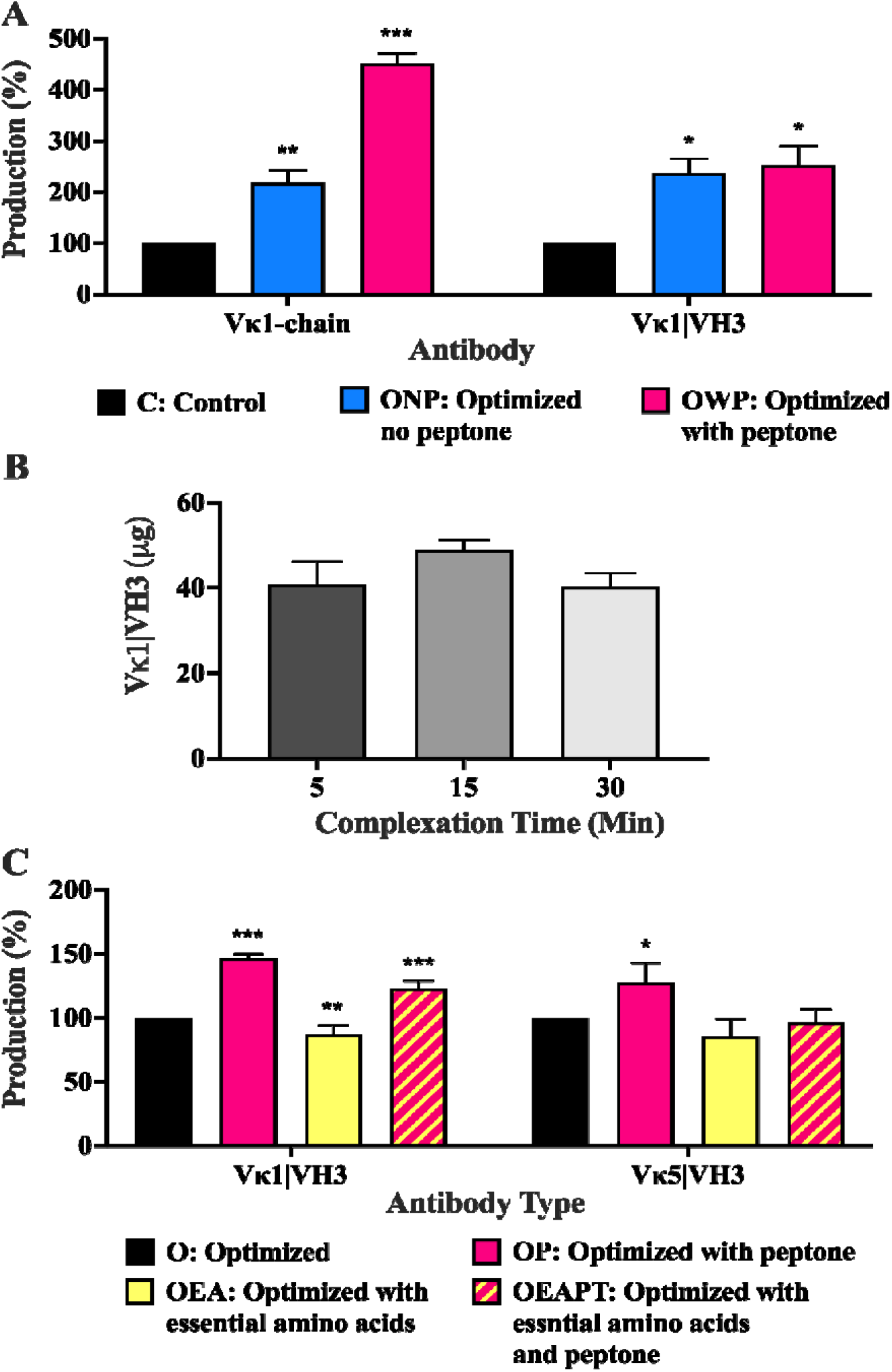
Production of recombinant Pertuzumab in 2mL transfections of (A) Light chain (Vκ1) and whole Vκ1|VH3 antibody among standard, optimised and with peptone supplementation conditions; (B) Whole Vκ1|VH3 Pertuzumab production at optimised parameters with PEI-DNA complex time variations; (C) Vκ1|VH3 and Vκ5|VH3 variants of recombinant Pertuzumab among optimised parameters with peptone and/or essential amino acids supplements. Black coloured bars in A and C represent the control to which the respective experiment parameters were compared by one-way ANOVA Dunnett’s post-hoc test. *, **. *** denote significance at *p* < 0.05, *p* < 0.01 and *p* < 0.001, respectively. (The experiments were carried out in at least three independent replicates where the total protein values are provided in Supplementary Table 3).

One-way ANOVA showed no significant difference (*p* > 0.05) in total whole antibody produced from initial 2 mL cultures with PEI-DNA complexation incubation times of 5, 15, and 30 minutes in **Figure 4B**. Following on from our previous work where 3.5 mg/ml of essential amino acids (EA) showed improvements in Pertuzumab antibody production (28), EA supplementation was added to our evaluation where we found an unexpected significant decrease (*p* < 0.05) of 13 % by Dunnett’s test, from an optimised control of 44 to 38 μg. (**Figure 4C** and Supplementary Table 3). Dual supplementations of both peptone and EA gave a significant increase (*p* < 0.05) of 23 % to 54 μg, with peptone alone giving a significantly higher (*p* < 0.05) increase of 47 % to 65 μg (**Figure 4C** and Supplementary Table 3). Comparing the single and dual addition of peptone and EA to boost light chain limiting antibodies (Pertuzumab Vκ5|VH3), we found peptone alone to significantly improve (*p* < 0.05) production by 28 % but not for other supplements. The average total protein data with accompanying standard error are shown in Supplementary Table 3.

## DISCUSSION

The cost-effective and efficient scalable production of recombinant monoclonal antibodies is key to the commercial success of both therapeutic and diagnostic antibodies. For this purpose, we set out to optimise transfection and culture parameters with relevance beyond antibody production to that of recombinant proteins using mammalian cell culture. Whole recombinant antibody production is affected by many factors such as the constant region (30) and CDRs of the Vκ and VH chains (29), their pairings (29), deletions in the Vκ (34), the signal peptides and amino acid usage (28). Given the importance of light chains for whole antibody production (26, 27) and to mitigate the possible lopsided effect of dual plasmid transfection, we performed single parameter optimisations on Pertuzumab light chain production. Thereafter, we applied the combined optimised parameters to produce the corresponding Pertuzumab.

Counter-intuitively to the assumption where higher cell seeding density could increase antibody production (39), we did not find significant differences in light chain production levels over the range of tested seeding densities (**Figure 1A**), hinting at the influence of other factors. In fact, high seeding cell densities significantly decreased production, which was consistent with previous literature (25), likely due to the reduction in transfection efficiency. It is important to note that the two cited papers here tested higher cell densities as suspension cultures while we used adherent HEK293E. For this, we found our typical usage of adherent cells in the range of 2 × 10^5^ cells/mL in 2 mL transfection within 6-well plates to be suitable as a baseline for further optimisations. Notably, there would be expected differences between suspension and adherent cell cultures.

With linear PEI previously reported to be a cost-effective (20), scalable (40), and efficient (41, 42) transfection reagent, we managed to obtain a peak light chain titre using up to a PEI concentration of 4 μg/mL with diminishing returns at higher concentrations. This result supports previous findings that increasing PEI to a certain peak improves protein expression but declines thereafter (25, 43). While increasing PEI can improve transfection efficiency, higher levels can be toxic, resulting in cell death, poorer transfection efficiencies and decreased recombinant protein production (12, 25). Although Dunnett’s test did not show a significant increase in light chain production between our control of 2 μg/mL and 4 μg/mL (**Figure 1B**), the t-test showed a significant increase of 45 %, pointing 4 μg/mL as optimal. Interestingly, DNA concentration as low as 0.25 μg/mL augmented light chain production (**Figure 1C**) and was thus incorporated into the optimised parameters. High amounts of DNA were likely to decrease production due to cellular toxicity associated with higher phosphoric acid levels (12).

Next, we optimised PEI-DNA complexation incubation conditions finding good agreement with other studies (12, 24, 44) that longer complexation incubation decreased protein production (**Figure 1D**). This is likely explained by the increase in PEI-DNA complex size with time (45, 46), reducing cell penetration and lowering transfection efficiency. From these findings, we reduced our complexation incubation time to five minutes. With temperature affecting PEI and DNA complexation speed, we also investigated the incubation temperatures, finding 18 °C to be optimal. Higher temperatures reduced light chain production (**Figure 1E**) possibly due to larger complex sizes being formed that decreased entry into cells (12). While these effects may not be the same for co-transfections of more than one type of plasmid, smaller complexes are likely to improve transfection efficiency for single plasmid transfections.

Further examining PEI-DNA complexation volume where complexation typically occurs in a volume of serum free medium (47) at 10 % of the total medium volume (20). Our results (**Figure 1F**) were in agreement with a previous study finding no significant differences between 5 and 10 % (12), and even at higher serum free medium percentages up to 25 %. From this, we retained the recommended 10 %. Since the serum free medium percentage had effects on light chain production, there were hints that reducing complexation volume could increase the proximity of PEI and DNA by enhancing its complexation to form larger complexes that could lead to poorer transfection efficiency (12).

As PEI-DNA complex is critical in the transfection process, we explored certain lab-culture practices with regards to complexation under light to explore if nucleic acid photodegradation (35) may affect transfection efficiency. Given that our results (**Figure 1 G**) show no significant difference in the light chain production, we ruled out photodegradation as a possible operational concern.

Turning to optimise culture conditions, we studied the effects of medium replacements, serum deprivation, culture in the presence of G418 antibiotic, culture incubation temperature, culture medium volume, post-transfection harvest time and supplements. Interestingly, we did not find medium replacements to cause any significant effects by one-way ANOVA, even though the t-test showed a significant increase in light chain production with fresh medium replacement prior to transfection (**Figure 2A**). This could be due to the transfection efficiency improving with waste product removal (7, 19). While such a practice is reasonable in laboratories, such a need could be costly and laborious (7) for larger-scale conditions. Interestingly, we found that medium replacements after transfection, conventionally performed to reduce transfection reagent toxicity unnecessary, for it reduced light chain production (**Figure 2A**).

With serum deprivation reported to improve transfection efficiency in some cell lines for purposes other than recombinant antibody production (14), we examined if serum deprivation could improve transfection efficiency and enhance light chain production. We found a non-significant decline in light chain production (**Figure 2B**) for the transfected HEK293E cells in agreement with a previous study (37). This could be due to the limited availability of specific essential amino acids from serum deprivation resulting in reduced antibody production (28).

Since G418 antibiotic is a requirement for HEK293E cells maintenance, we investigated if it could potentially increase metabolic stress and affect recombinant protein production after transient transfection (15). Since our results (**Figure 2C**) showed no significant difference in the light chain production with its addition, large-scale protein production can be performed with or without G418.

Based on the assumption that growth rate retardation would retain higher cell viability, lower nutrient consumption rate and reduce waste accumulation, amongst better mRNA stability, improved protein folding, and fidelity of post-translational events (48), cell culture incubation temperature was investigated. Notably, the literature in this area was inconclusive. Some were reportedly better (49), having no effect (50) or decreasing production (51) at lower temperatures such as 32 °C, with only one study on HEK293S suspension cells finding better production of non-antibody recombinant proteins at lower temperatures (13). Controlling for different evaporation rates by performing normalisations using total protein data from 2 mL transfections, we found lower temperatures detrimental to light chain production (**Figure 2D**). An extended experiment using larger 40 mL T175 transfections with a panel of Pertuzumab and Trastuzumab whole antibody variants showed no clear trends for production (Supplementary Figure 1A and B). Thus, we continued using the optimal growth temperature of 37 °C.

On the reasonable assumption that the culture medium would provide the necessary nutrients for protein production, we investigated if increasing the culture medium would overcome nutrition bottlenecks and improve light chain production. Since different culture volumes were used, total protein production data from experimental conditions were normalised to the control. Dunnett’s test showed significantly lower light chain production with the usage of 1 mL cultures, but there were no significant increases despite a 40 % increase with the use of 5 mL cultures (**Figure 2F**). Apart from the significantly lower light chain production, culture volumes below 2 mL are generally not recommended as evaporation becomes an issue. On the contrary, volume increases in 6-well plates with possible spillage and culture medium cost are not practical in larger-scale productions.

Following this, we explored post-transfection harvest, finding a direct relationship between post-transfection harvest time and light chain production which plateaued on day 14. Depending on the downstream application and the quantity of antibody desired, our results suggest that culture supernatant can be reasonably harvested anytime between day 3 to 21 post-transfection (**Figure 2F**), preferably between days 5 to 14 (52, 53). It should also be noted that longer culture times could lead to the accumulation of host-cell proteins, possibly confounding subsequent purification processes. This consideration may also be useful for proteins with poor stability where prolonged culture conditions can cause degradation.

Exploring the addition of supplements, peptone (25, 36–38), tryptone (38), casamino acids (54), yeast extract (55), casein (56), and skim milk (57) were tested. Given that the high amino acid content in the supplements could lead to toxic accumulation of ammonia culminating in cellular toxicity (36), we tested them at 1 mg/mL. Peptone was found to significantly boost light chain production in agreement with previous reports (25, 36–38). However, light chain production was not augmented by other supplements (**Figure 3A**). Determining the optimal time of supplementation, addition of 1 mg/mL peptone on either post-transfection days 5, 6 or 7 were best in boosting production (**Figure 3B**). Repeated additions of supplements did not yield better productions (**data not shown**), possibly due to toxic accumulation of ammonia (36).

Combining the optimised parameters, we improved Pertuzumab light chain production in 2 mL transfections by 217 % to 536 μg, which was further enhanced ~2-fold to 452 % reaching 1032 μg with peptone supplementation (**Figure 4A** and Supplementary Table 3). Applied to whole Pertuzumab antibody in 2 mL transfections, a similar increase of 236 % from 22 μg to 49 μg was observed, with peptone addition further significantly augmenting production to 252 % reaching 51 μg (**Figure 4A** and Supplementary Table 3). Hypothesising that dual plasmid transfection might require a longer PEI-DNA complexation time, we attempted to optimise it for whole antibody production but did not observe significant differences (**Figure 4B**). From a possible lead in our previous work (28) where 3.5 mg/mL EA supplementation improved whole antibody production, we explored both single and dual supplementations of EA and peptone. We found an unexpected significant whole antibody production decrease upon EA addition which was rescued by the addition of peptone (**Figure 4C**) for the Pertuzumab antibody.

Considering that some whole antibody productions are limited by the light chain (i.e. Pertuzumab Vκ5) from our previous work (29) and that peptone appeared to benefit light chain production in this study, we performed similar experiments using Pertuzumab Vκ5|VH3 whole antibodies for validation. We were able to significantly boost the whole Vκ5|VH3 antibodies production (**Figure 4C**) by 28 % from 2.4 to 3.0 μg in 2 mL transfections (Supplementary Table 3) supporting the use of peptone supplements for antibodies limited by light chains in transient production. Thereby, peptone could be added for light chain limiting antibody production given its benefits and lack of detrimental effects.

## CONCLUSION

Through a detailed systematic selection of antibodies in a holistic (1, 31) manner, the selection of previously published factors of heavy chain constant (30), Vκ-VH pairings (29), amino acid usage with signal peptide selection (28), antibody production could be more efficient, mitigating commercial failures due to production costs. Here our optimisations of the parameters led to a ~2-fold increase of light chain only and whole antibody production from HEK293E using 25 kDa linear PEI in a cost-effective and operationally convenient manner. With peptone supplements, light chain increased by ~4-fold while whole antibody increased to a lesser extent, with most benefits found for antibodies limited by their light chains in production. These findings have potential applications to other mammalian-based recombinant protein production.

## DATA AVAILABILITY STATEMENT

The data underlying this article will be shared at reasonable request to the corresponding author.

## AUTHOR CONTRIBUTIONS

Conceptualisation, W.-L.L and S.K.-E.G.; methodology, W.-L.L. and Z.S.-L.H.; formal analysis, W.-L.L., Z.S.-L.H.; S.K.-E.G; investigation, W.-L.L. and Z.S.-L.H.; validation, D.W.-S.K., J.Y.Y., writing—original draft preparation, Z.S.-L.H; writing—review and editing, D.W.-S.K., J.Y.Y., S.K.-E.G., W.-L.L. and Z.S.-L.H; visualisation, S.K.-E.G., W.- L.L. and Z.S.-L.H; supervision, S.K.-E.G. and W.-L.L; funding acquisition, S.K.-E.G. All authors have read and agreed to the published version of the manuscript.

## FUNDING STATEMENT

This work was initially supported by the Joint Council Office, Agency for Science, Technology, and Research, Singapore, under Grant number JCO1334i00050 and later by the National Research Foundation (NRF) Singapore grant to Experimental Drug Development Centre (EDDC) for the platform AMDED, acquired by S.K.-E.G.

## ACKNOWLEDGEMENTS

The authors would also like to thank Jun-Jie Poh for his T175 flasks transfection data contribution.

## CONFLICT OF INTEREST STATEMENT

The authors declare no conflict of interest.

## ETHICS AND CONSENT STATEMENT

Consent was not required for this study

## ANIMAL RESEARCH STATEMENT

This is not applicable to this study.

## Notes

### Competing Interest Statement

The authors have declared no competing interest.

### Summary of Updates

Updated manuscript.

