## Supplementary for "Augmenting Recombinant Antibody Production in HEK293E Cells: Optimising Transfection and Culture Parameters"

**Supplementary Table 1**: Average Pertuzumab light chain concentrations from at least three independent replicates attained in 6-well plates, 2 mL transfections for the investigation of transfection or culture parameters.

| Parameters | Conditions | Average Protein Titre (μg/mL) | Standard Error |
|---|---|---|---|
| Cell Density (cells/mL) | 1 x $10^5$ | 174.6 | 10.2 |
| | 2 x $10^5$ (Control) | 177.1 | 18.3 |
| | 4 x $10^5$ | 185.6 | 15.5 |
| | 6 x $10^5$ | 149.3 | 33.9 |
| | 8 x $10^5$ | 92.1 | 29.6 |
| | 10 x $10^5$ | 69.7 | 22.0 |
| PEI Concentration (μg/mL) | 1 | 21.7 | 5.4 |
| | 2 (Control) | 139.5 | 22.8 |
| | 4 | 206.0 | 45.4 |
| | 6 | 172.6 | 74.9 |
| | 8 | 92.7 | 50.1 |
| | 10 | 57.3 | 30.6 |
| DNA Concentration (μg/mL) | 0.25 | 398.6 | 49.1 |
| | 0.50 | 323.2 | 23.4 |
| | 0.75 | 235.9 | 5.8 |
| | 1.00 (Control) | 208.2 | 15.7 |
| | 1.25 | 166.1 | 18.1 |
| | 1.50 | 67.1 | 10.9 |
| | 2.00 | 64.2 | 3.1 |
| Complexation Time (min) | 5 | 271.1 | 69.6 |
| | 10 | 233.2 | 49.7 |
| | 20 | 222.9 | 58.0 |
| | 30 (Control) | 164.6 | 43.4 |
| | 40 | 146.7 | 56.8 |
| | 60 | 108.6 | 42.2 |
| | 90 | 86.1 | 35.3 |
| | 120 | 53.6 | 16.1 |
| Complexation Temperature (°C) | 18 (Control) | 163.9 | 65.8 |
| | 25 | 122.3 | 50.2 |
| | 32 | 139.6 | 58.7 |
| | 37 | 128.0 | 52.4 |
| | 40 | 104.8 | 47.7 |
| | 45 | 98.3 | 46.5 |
| Complexation Volume (%) | 1 | 30.0 | 12.7 |
| | 5 | 136.6 | 33.9 |
| | 10 (Control) | 213.6 | 55.8 |
| | 15 | 254.8 | 52.9 |

| | | | |
|---|---|---|---|
| | 20 | 255.3 | 47.6 |
| | 25 | 272.5 | 73.5 |
| Complexation Environment | Dark (Control) | 158.4 | 26.2 |
| | Light | 182.8 | 6.6 |
| Medium Replacement | No medium replacement (Control) | 226.1 | 74.8 |
| | Before transfection | 319.7 | 120.4 |
| | Before and after transfection | 250.4 | 84.9 |
| | After transfection | 235.0 | 126.9 |
| Serum Deprivation | No serum deprivation (Control) | 226.1 | 74.8 |
| | Serum deprivation | 175.4 | 63.3 |
| | Serum deprivation, media replacement after transfection | 145.3 | 51.0 |
| G418 Antibiotic | Present | 209.3 | 14.8 |
| | Absent (Control) | 209.4 | 19.6 |

**Supplementary Table 2:** Average amount of Pertuzumab light chain produced from at least three independent replicates attained in 6-well plates, 2 mL transfections for the investigation of cell culture parameters taking into account evaporation over time.

| Parameters | Conditions | Total Protein (μg) | Standard Error |
|---|---|---|---|
| Culture Temperature (°C) | 32 | 120.2 | 12.4 |
| | 37 (Control) | 231.1 | 7.2 |
| Medium Volume (mL) | 1.0 | 148.2 | 71.8 |
| | 1.5 | 293.9 | 51.5 |
| | 2.0 (Control) | 299.9 | 12.3 |
| | 2.5 | 314.1 | 8.8 |
| | 3.0 | 293.7 | 30.8 |
| | 3.5 | 332.2 | 48.1 |
| | 4.0 | 361.9 | 56.9 |
| | 4.5 | 345.8 | 36.2 |
| | 5.0 | 413.4 | 32.4 |
| Post-transfection Harvest (Day) | 3 | 80.6 | 36.7 |
| | 6 | 251.6 | 105.9 |
| | 9 | 371.6 | 176.0 |
| | 12 | 462.6 | 202.2 |
| | 14 (Control) | 522.1 | 189.3 |
| | 18 | 532.3 | 189.2 |
| | 21 | 525.3 | 160.8 |
| Supplements | No supplement (Control) | 277.9 | 49.3 |
| | Casaminoacids | 296.0 | 73.2 |
| | Casein | 267.9 | 56.9 |
| | Peptone | 350.6 | 51.1 |
| | Tryptone | 261.8 | 53.6 |
| | Skim milk | 281.3 | 58.7 |
| | Yeast Extract | 336.1 | 72.2 |

**Supplementary Table 3:** Average amount of Pertuzumab light chain and whole antibody produced from at least three independent replicates attained in 6-well plates, 2 mL transfections for the investigation of the optimized parameters taking into account evaporation over time.

| Experimental Set | Antibody Type | Conditions | Total Protein (µg) | Standard Error |
|---|---|---|---|---|
| 1 | Vκ1 light chain | Control protocol | 265.1 | 78.1 |
| | | Optimised protocol | 535.8 | 122.2 |
| 2 | Vκ1 light chain | Control protocol | 228.0 | 5.1 |
| | | Optimised protocol with peptone | 1032 | 64.9 |
| 3 | Vκ1 light\|VH3 heavy chains | Control protocol | 21.5 | 3.8 |
| | | Optimised protocol | 48.5 | 2.4 |
| | | Optimised protocol with peptone | 51.1 | 0.6 |
| 4 | Vκ1 light \|VH3 heavy chains | Optimised protocol | 44.2 | 2.2 |
| | | Optimised protocol with peptone | 64.7 | 2.8 |
| | | Optimised protocol with essential amino acids | 38.4 | 1.7 |
| | | Optimised protocol with essential amino acids and peptone | 54.2 | 1.4 |
| 5 | Vκ5 light\|VH3 heavy chains | Optimised protocol | 2.4 | 0.1 |
| | | Optimised protocol with peptone | 3.0 | 0.2 |
| | | Optimised protocol with essential amino acids | 2.0 | 0.1 |
| | | Optimised protocol with essential amino acids and peptone | 2.3 | 0.1 |

**Analysis of incubation temperatures 32 °C and 37 °C on antibody production**

To establish the effects of post-transfection incubation temperatures on antibody production, we tested a variety of VH and Vκ paired Pertuzumab and Trastuzumab CDR-grafted antibodies incubated at either 32 °C or 37 °C as shown below. Generally, no clear trends were established, suggesting the need for tailored optimisation of specific antibodies when upscaling.

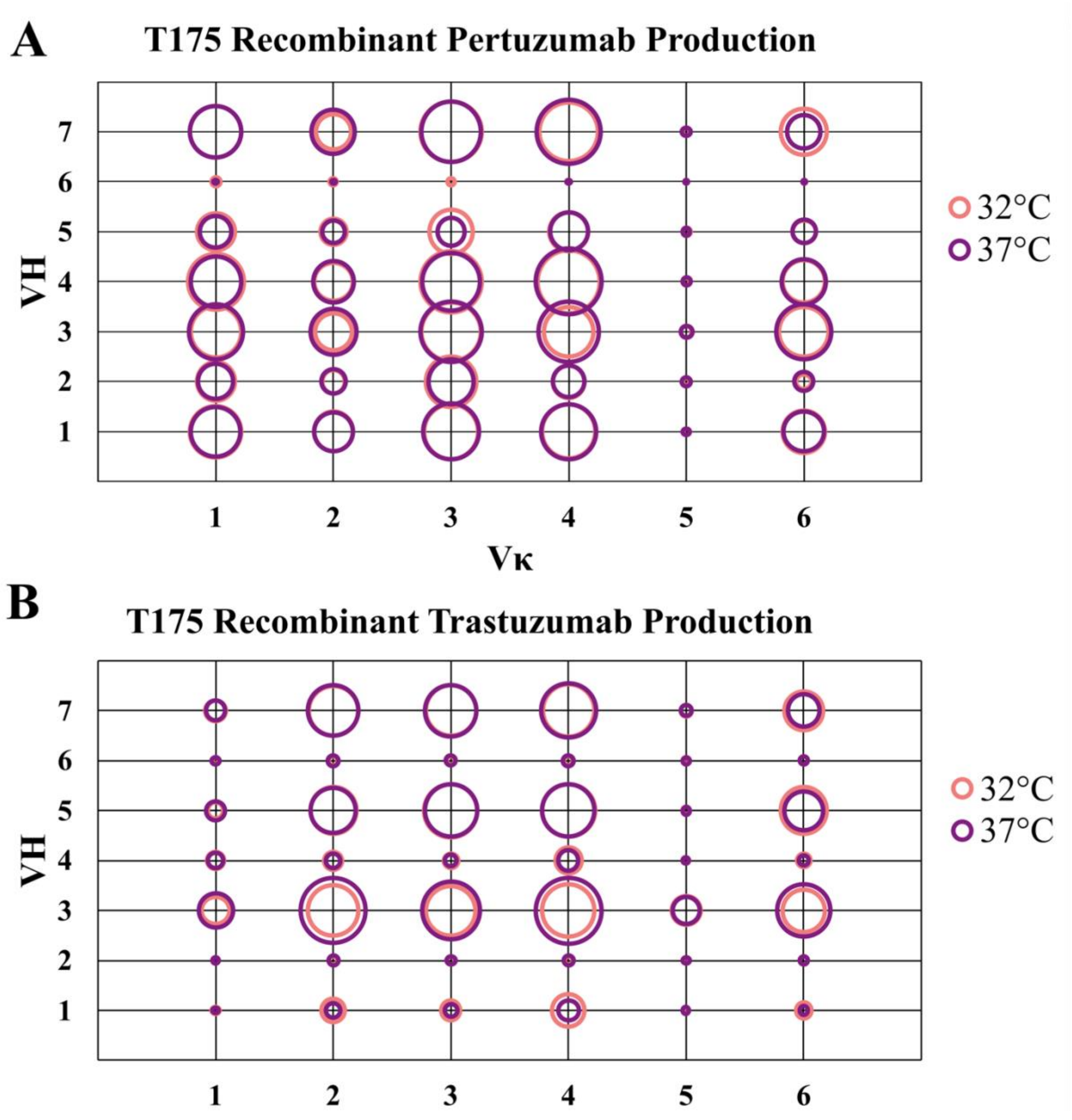

**Supplementary Figure 1:** Bubble chart representations of T175 flasks, 40 mL transfections of a panel of recombinant Pertuzumab (A) and Trastuzumab (B) variants (VH and Vκ) previously described (1, 2). Each circle in the respective combinations shows the production level of the antibody comparing culture incubation temperatures of 32 °C and 37 °C.
